# RNase III in *Salmonella* Enteritidis enhances bacterial virulence by reducing host immune responses

**DOI:** 10.1101/2024.09.04.611193

**Authors:** Bill Kwan-wai Chan, Yingxue Li, Hongyuhang Ni, Edward Wai-chi Chan, Xin Deng, Linfeng Huang, Sheng Chen

## Abstract

*Salmonella* is an important foodborne pathogen which comprises strains that exhibit varied virulence phenotypes and the capability of causing invasive human infection. In this study, the gene expression profile of foodborne and clinical *Salmonella* strains that exhibit high- and low-level virulence was investigated, with results showing that the expression level of a number of genes, including the *rnc* gene which encodes the RNase III ribonuclease, were exceptionally high in the high virulence strains. Investigation of the role of *rnc* in mediating expression of virulence phenotypes in *Salmonella* showed that the product of this gene could enhance expression of the superoxide dismutase SodA, which is an essential determinant of survival fitness of *Salmonella* under the oxidative stress elicited by the host immunity. On the other hand, we also discovered that the double-stranded RNA (dsRNA) released from *Salmonella* could trigger immune response of the host, and that the high-level expression of the *rnc* gene enabled *Salmonella* to evade the host immunity by reducing the amount of dsRNA accumulated in the bacterial cell. These findings provide insightful understanding of the regulation of *Salmonella* virulence and facilitate development of novel antimicrobial treatments through suppression of virulence expression and survival fitness of this important pathogen.

## INTRODUCTION

*Salmonella enterica* is one of the most important foodborne pathogens, causing over one million cases of infection in the United States annually (1). Non-typhoidal *Salmonella* account for the majority of infections. To date, about 2500 serovars of *S. enterica* have been recognized, two of which, namely Typhimurium and Enteritidis, are responsible for most cases of non-typhoidal Salmonellosis (2). Non-typhoidal *Salmonella* infections are mostly associated with self-limiting diarrhea. However, invasion of the pathogen into normally sterile sites, including bloodstream and meninges, is possible, resulting in focal infections (3). Invasiveness of non-typhoidal *S. enterica* into various body sites has been observed worldwide, however, the mechanisms underlying this invasion process remain poorly understood.

We have observed that *Salmonella* strains collected from food and clinical samples exhibited a highly diverse range of invasion and survival ability on RAW264.7 macrophage cells (**Figure S1**). When subjected to comprehensive genetic analysis and different virulotyping assays to investigate the relationships between the genetic profiles of the test strains and their virulence level, we found that genetically identical strains might exhibit highly variable virulence levels, and that the molecular basis of such differences was due to variation in expression levels of specific gene clusters. Among these over-expressed gene clusters associated with higher virulence, we found that the *rnc*-*era*-*recO* operon was up-regulated in the highly virulent *Salmonella* isolates (4). The *rnc*-*era*-*recO* operon has been identified in various types of bacteria (5). In this operon, the *rnc* gene encodes the RNase III ribonuclease, which specifically cleaves double stranded RNA (dsRNA), resulting in the formation of a two-nucleotide 3’ overhang at each end of the cleaved dsRNA (6). It was reported that the *rnc* gene in *E. coli* played an important role in regulation of protein synthesis (7) and degradation of structured RNAs and dsRNAs formed by overlapping sense and antisense RNAs (8). However, the exact function of the *rnc* gene in regulating *Salmonella* virulence is unknown.

In this study, we investigated the role of *rnc* gene in mediating virulence phenotype in *Salmonella.* Two potential functions of RNase III were tested firstly in regulating gene expression and secondly in mediating enhanced resistance to the host immune response. We found that the *rnc* gene is the key determinant of virulence in *Salmonella*. Findings in this work could facilitate development of novel strategies to suppress the virulence level of *Salmonella*.

## RESULTS

### The rnc gene was over-expressed in high virulence strains

A total six *S.* Enteritidis strains were recovered from food and clinical samples. Their virulence phenotypes were characterized by determining the internalization and intracellular replication rate of these strains in macrophage RAW264.7 (**Table 1**). The macrophage internalization and intracellular replication rate of the food isolates (654, 2992 and 3046) ranged from 0.0075 to 0.0152, and 0.0370 to 1.0150 respectively. Those of the clinical isolates (SE 12-5, SE 11-72 and SE 09-1889) ranged from 0.2189 to 0.2925, and 4.4744 to 15.3199 respectively. The results indicated that the food isolates generally exhibited a lower level of virulence when compared to the clinical isolates. However, it should be noted that strain 654 exhibited exceptionally high virulence among the three food isolates tested, suggesting that the high-virulence *Salmonella* strains exist in the natural environment.

**Table 1.** The macrophage internalization and replication rate of six *S.* Enteritidis strains.

| Strains | Source | Location | Year | Internalization rate | Replication rate |
| --- | --- | --- | --- | --- | --- |
| 2992 | food | China CDC | 2012 | 0.0152 | 0.7934 |
| 3046 | food | China CDC | 2012 | 0.0135 | 0.0370 |
| 654 | food | China CDC | 2012 | 0.0075 | 1.0150 |
| SE 12-5 | stool | CUHK | 2012 | 0.2189 | 7.6038 |
| SE 11-72 | stool | CUHK | 2011 | 0.2925 | 4.4744 |
| SE 09-1889 | blood | CUHK | 2009 | 0.2228 | 15.3199 |

All tested strains were further subjected to RNA sequencing and the results showing that specific genes were over-expressed in the high virulence isolates recovered from the clinical samples as well as the food isolate 654. These highly expressed genes include a number of virulence determinants (e.g. Type-3 secretion system and fimbriae protein-encoding genes), transcription regulators (e.g. *araC* family transcription regulators and the *rnc-era-recO* operon) and various genes involved in cellular functions related to metabolism. Among those genetic elements, the *rnc* gene which encodes RNase III was found to be over-expressed in all clinical *S*. Enteritidis isolates (**Figure 1a**). Observation of high intracellular replication rate of the tested isolates (**Table 1**) was highly consistent with high expression level of the *rnc* gene. These findings suggest that the *rnc* gene may play a role in mediating expression of virulence in *S*. Enteritidis. However, the RNA sequencing data couldn’t reveal the genetic basis of the significant differences in *rnc* gene expression level among the high and low virulence strains at this stage.

**Figure 1.**
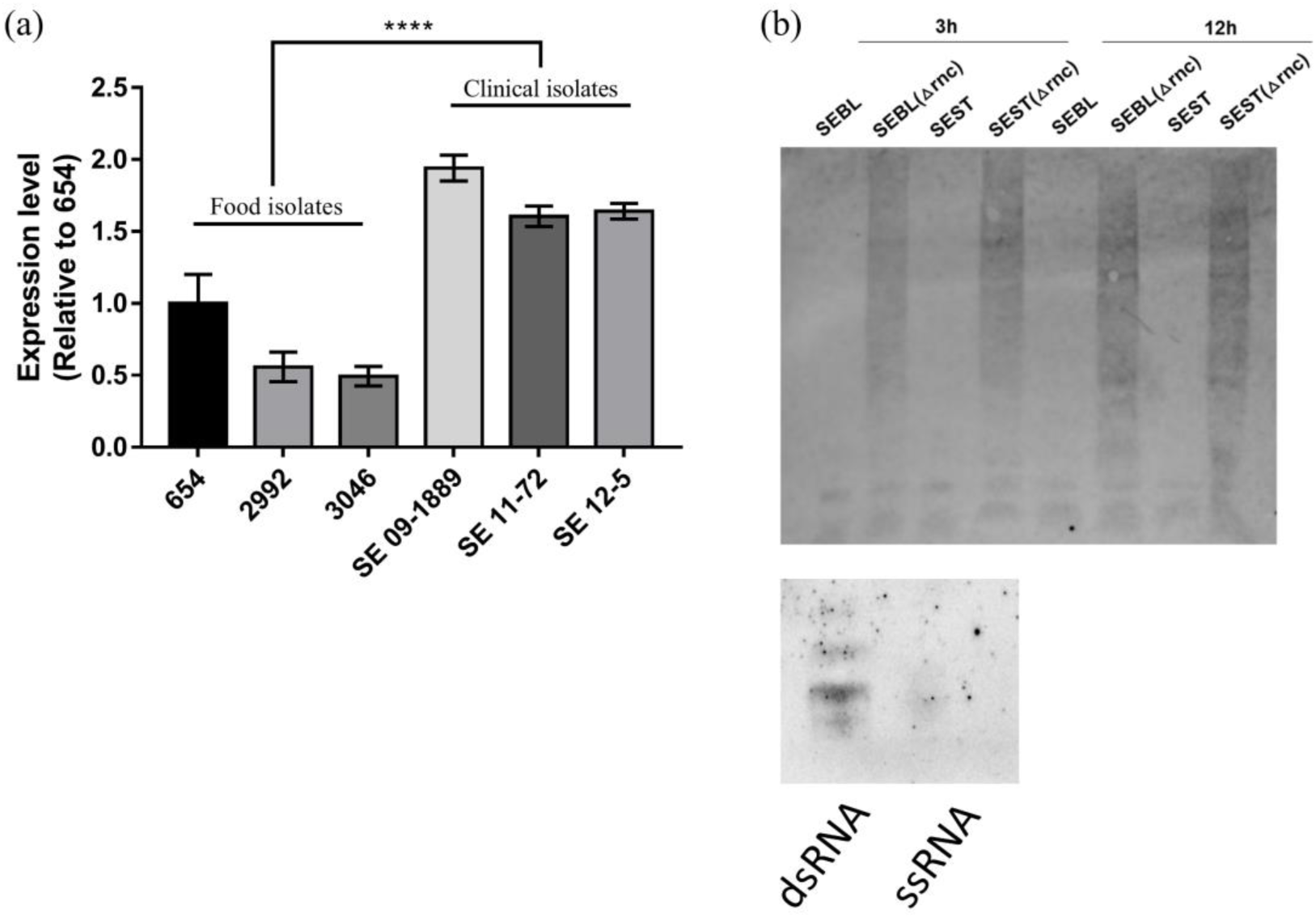
RNase III was over-expressed in high virulence strains and was responsible for degradation of dsRNA. (a) Expression level of the *rnc* gene in six *S.* Enteritidis isolates. (b) Two clinical isolates (SEBL, SEST), and their corresponding *rnc* knockout mutants (SEBLΔ*rnc*, SESTΔ*rnc*) were cultured for 3h and 12h. Their total RNAs were extracted and the dsRNA was detected using J2 antibody. Same amount of total RNA from each sample were loaded. The specificity of J2 antibody on dsRNA was checked by using commercial dsRNA and ssRNA ladder.

Since the function of *rnc* gene product RNase III ribonuclease involves degradation of dsRNA, we hypothesized that the alteration of virulence level in *Salmonella rnc* mutant might be due to intracellular accumulation of dsRNA. In order to test whether dsRNA was accumulated in the *rnc* knockout mutant, immunoblotting against dsRNA was performed. The *Salmonella* strains, SEBL, SEST, SEBLΔ*rnc* and SESTΔ*rnc* were tested in this experiment. Upon being cultured for 3 h and 12 h, total RNAs were extracted and the dsRNA was detected using the dsRNA specific J2 antibody (**Figure 1b**). The results showed that dsRNA was not detectable in both 3 and 12 h culture samples from the wildtype SEBL and SEST strains. However, dsRNA could be detected in the 3 h culture samples of the SEBLΔ*rnc* and SESTΔ*rnc* mutant strains and the quantity of dsRNA was much higher in the 12 h culture samples. These findings indicate that RNase III was responsible for the degradation of intracellular dsRNAs in *Salmonella* and that the low level of dsRNA correlated with high level virulence in wildtype *Salmonella*.

### The rnc gene was essential for virulence phenotype in Salmonella

To further confirm the role of the *rnc* gene product in regulating virulence expression in *Salmonella*, the *rnc* gene was deleted from the high virulence foodborne isolate 654 to create the 654Δ*rnc* mutant, which was then subjected to internalization and replication assay in RAW264.7 macrophage cell line (**Figure 2a, b**). The result showed that both internalization and intracellular replication rate of *S*. Enteritidis 654 in macrophage cells were reduced by up to 80% when the *rnc* gene was deleted from the genome (**Figure 2b**). Consistently, the internalization and intracellular replication rates of the 654Δ*rnc*/p-*rnc* strain, which was created by transformation of a *rnc* gene-bearing plasmid into 654Δ*rnc* mutant to complement the lack of *rnc* in the mutant, were found to have reverted back to the similar virulence level as *S*. Enteritidis 654. These findings confirmed that *rnc* gene was important for *Salmonella* survival inside macrophage cells and the overall infection ability of this important pathogen. In summary, *rnc* gene is a key virulence determinant of *Salmonella*.

**Figure 2.**
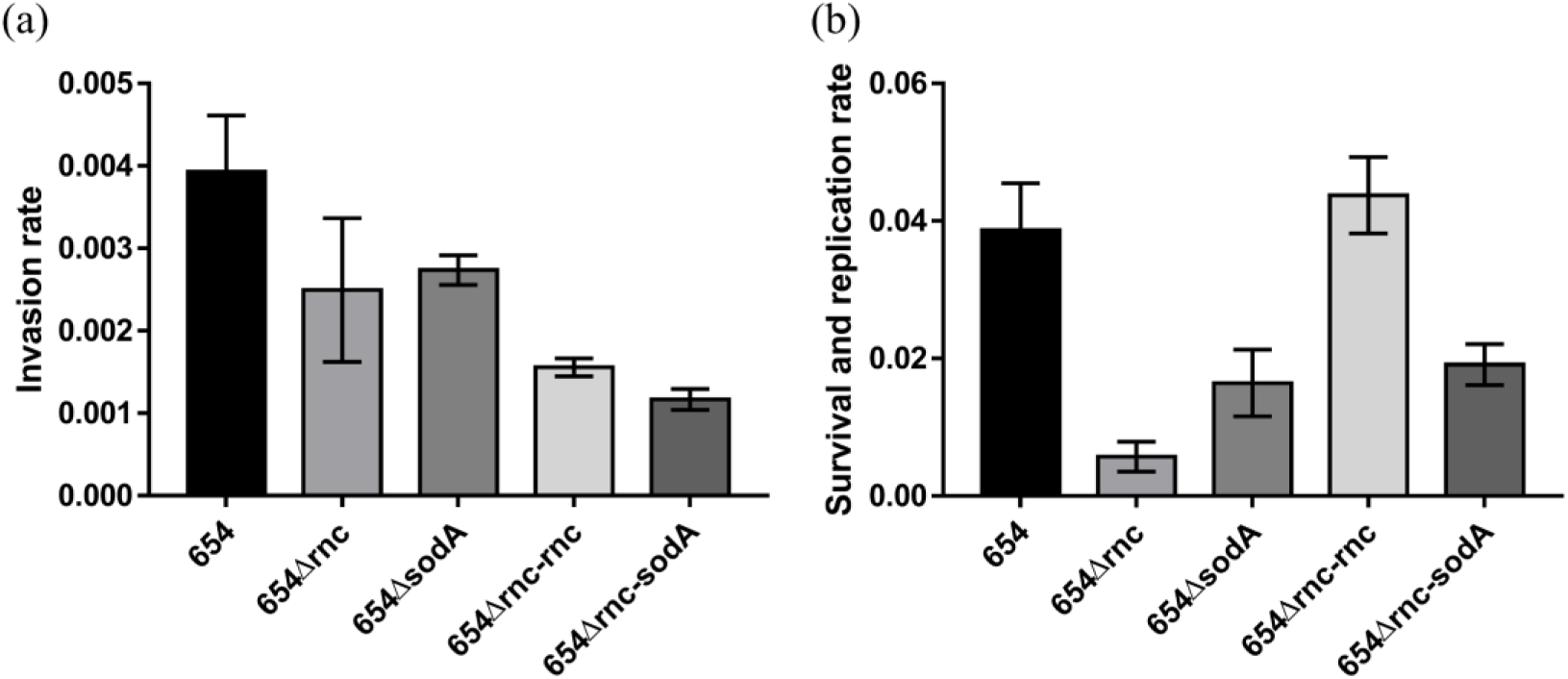
Virulence assays of *Salmonella* strains carrying different constructs. (a) RAW264.7 invasion rate of strains carrying different constructs. (b) RAW264.7 intracellular survival and replication rate.

### Double-stranded RNA in Salmonella could induce immune response in the host

Detection of double-stranded RNA is an important feature of innate immune system in many organisms, especially for defense against viruses which frequently employed double-stranded RNA as their genetic materials (9). A known function of the gene product of *rnc* in bacteria is to digest double-stranded RNA (dsRNA) and we showed *Salmonella rnc* mutants that contained higher levels of dsRNAs (**Figure 1b)**. We hypothesized the high expression of *rnc* gene in high virulence *Salmonella* strains could promote the elimination of dsRNA to minimize the activation of host immune response to dsRNA during infection. To investigate whether the host immune response could be triggered by dsRNA of *Salmonella*, total RNA extracted from *S*. *Enteritids* 654 strains carrying different constructs were used to transfect different mammalian cell lines, followed by monitoring the changes in immune factors. Expression levels of TNF-α (**Figure 3a)**, IL-1β (**Figure 3b**), MDA-5 (**Figure 3c**) and IFN-β (**Figure 3d**) could be detected in RAW264.7 macrophage cells; on the other hand, expression of IFN-β (**Figure 3e**), MDA-5 (**Figure 3f**), and RIG-I (**Figure 3g**) were detectable in Caco-2 colon epithelial cells. The results showed that all the immune related genes tested were inducible in both RAW264.7 and Caco-2 cells when they were transfected with total RNA extracted from strains bearing different constructs, suggesting that the RNA from *Salmonella* would stimulate the host immune response. The RAW264.7 cells transfected with total RNAs extracted from *S*. Enteritidis strain 654 and 654Δ*rnc*/p-*rnc* exhibited similar expression levels of TNF-α, IL-1β, MDA-5 and IFN-β (**Figure 3a-d**). However, cells transfected with total RNAs from 654Δ*rnc* exhibited significantly higher expression of immune factors; in particular, the level of expression of IL-1β and IFN-β was more than 50% higher in 654Δ*rnc* RNA transfected RAW264.7 cells (**Figure 3b,d**). The expression of IFN-β and RIG-I in Caco-2 cells transfected with total RNA extracted from strain 654Δ*rnc* was 4-fold that of the cells transfected with total RNA of strain 654 and 654Δ*rnc*/p-*rnc* (**Figure 3e, g**). The expression level of MDA-5 in Caco-2 cells was increased for 10-fold in 654Δ*rnc* RNA transfected cells (**Figure 3f**). These findings showed that total bacterial RNAs containing abundant dsRNAs from strain 654Δ*rnc* could induce strong innate immune responses in both RAW264.7 and Caco-2 cells.

**Figure 3.**
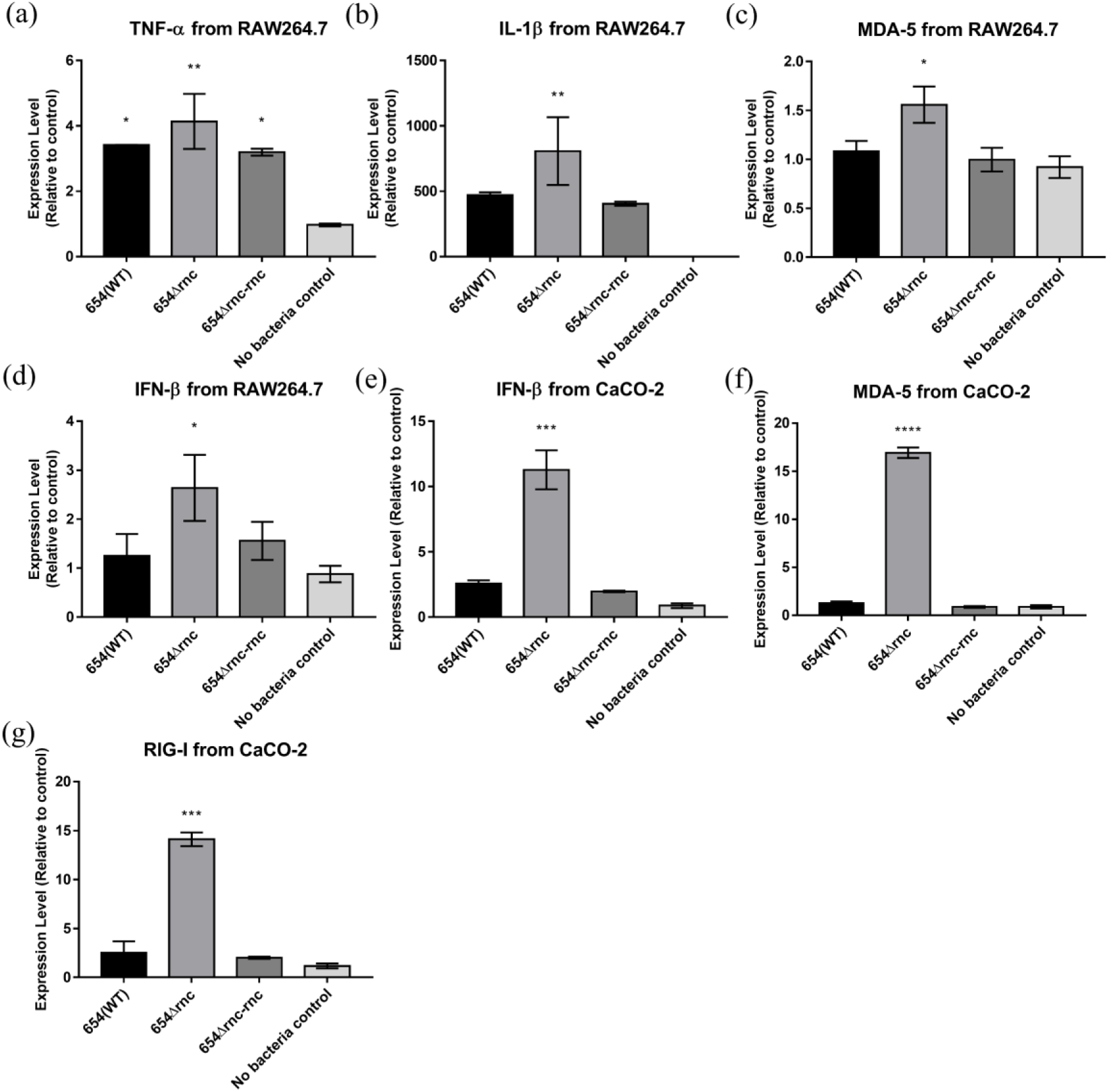
Expression of immune signals in macrophage RAW264.7 and colon epithelial cells Caco-2 upon being transfected by total RNA extracted from *Salmonella* strains carrying different constructs. RNA was transfected into the cell using Lipofectamine 2000 transfection reagent. (a) Expression of (a)TNF-α, (b)IL-1β, (c)MDA-5, (d)IFN-β in RAW264.7 cell line; Expression of (e)IFN-β, (f)MDA-5, (g)RIG-1 in Caco-2 cell line. The cells of no-bacteria control were treated with Lipofectamine 2000 transfection reagent without RNA.

To further confirm that the host immune responses were triggered by dsRNAs of *Salmonella*, total RNAs extracted from two clinical isolates, namely SEBL and SEST, and their corresponding *rnc* knockout mutants SEBLΔ*rnc* and SESTΔ*rnc*, were used to transfect the RAW264.7 macrophage cells and Caco-2 colon epithelial cells. Consistent with the above-described results from experiment involving the 654Δ*rnc* and 654Δ*rnc*/p-*rnc* strains, the mRNA expression level of IFN-β in RAW264.7 transfected with the total RNAs of SEBLΔ*rnc* and SESTΔ*rnc* was 100-fold and 200-fold higher than that of transfected with RNAs from wildtype SEBL and SEST, respectively (**Figure 4a**). Similarly, an around two-fold increase in mRNA expression of both MDA-5 and RIG-I was observed in cells transfected with total RNAs comparing SEBLΔ*rnc* with SEBL and comparing SESTΔ*rnc* with SEST, an around three-fold such increase was observed (**Figure 4b, c**). The expression level of IFN-β in Caco2 cells was also increased for 30-folds after transfection with total RNAs extracted comparing *rnc* mutants (SEBLΔ*rnc* or SESTΔ*rnc*) with wildtypes (SEBL and SEST).

**Figure 4.**
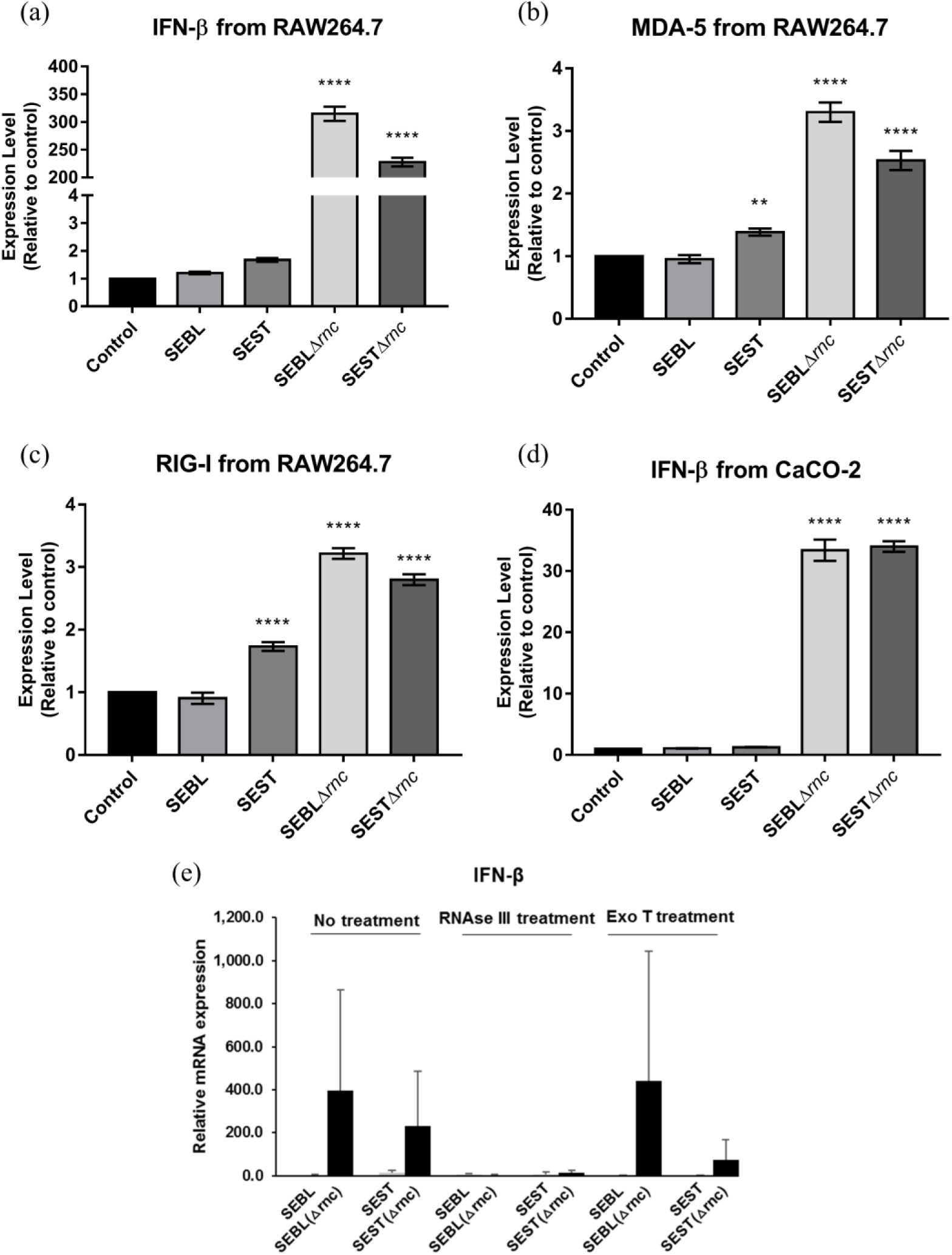
Expression of immune signals in macrophage RAW264.7 and colon epithelial cells Caco-2 upon being transfected by total RNA extracted from clinical *Salmonella* isolates. (a to d) RNAs extracted from strain SEBL and SEST, and their corresponding knockout mutants SEBLΔ*rnc* and SESTΔ*rnc* were transfected into two cell lines. The expression levels of immune signal were detected. (e)Total RNAs of the four *Salmonella* strains were extracted and treated with RNase III (dsRNA endonuclease) and exonuclease T (single-stranded RNA nuclease) respectively, followed by transfection into Caco-2 cell line to elicit different expression level of IFN-β. The cells treated with Lipofectamine 2000 only were used as control.

To further confirm that the increased immune responses in RAW264.7 and Caco-2 cells were triggered by the dsRNA instead of single-stranded RNA (ssRNA) from *Salmonella*, total RNAs from SEBL, SEBLΔ*rnc*, SEST and SESTΔ*rnc* were treated with dsRNA-specific RNase III and ssRNA-specific Exonuclease T respectively, followed by transfection into RAW264.7 cells and measurement of IFN-β expression (**Figure 4e**). The result showed that the expression levels of IFN-β in cells transfected with Rnase III-treated RNA samples from *rnc* mutants were reduced to the level of WT RNA. However, total RNAs from *rnc* mutants treated with exonuclease T could still induce high expression of *IFN-β* in RAW264.7 cells in a manner resembling the no-treatment group (**Figure 4e**). These findings strongly support that the induction of *IFN-β* expression in RAW264.7 cells was mainly due to dsRNA, not ssRNA of *Salmonella*.

In summary, these findings confirm that the total RNAs and particularly dsRNAs from *rnc* knockout mutants could stimulate significant immune responses in both RAW264.7 and Caco-2 cells. The expression of the *rnc* gene in WT resulted in reduced immune stimulatory effects of bacterial RNAs, thereby facilitating bacterial infection process.

### The rnc gene regulates expression of the superoxide dismutase SodA

The results of macrophage infection assays indicated that intracellular survival and replication of *S*. Enteritidis 654 in macrophages were regulated by the *rnc* gene product (**Figure 2**). Since the survival of *Salmonella* in macrophage was reported to be highly dependent on its ability of reactive oxygen species (ROS) detoxification, we further tested the ROS level in the six *Salmonella* isolates upon the treatment with hydrogen peroxide (**Figure 5a**) (10). A significantly lower level of ROS was detectable in the high virulence strains SE09-1889, SE11-72 and SE12-5 when compared to the low virulence strains 2992 and 3046, whereas intermediate level of ROS was recorded in strain 654, suggesting that the ROS level in *Salmonella* was inversely correlated with their virulence level as well as the expression level of *rnc* (**Figure 1**). Based on these observations, we hypothesize that RNase III expression may enhance the expression of superoxide dismutase in *Salmonella* and that this is one of the reasons why *rnc* is important for survival of *Salmonella* against the host defense systems. To test this hypothesis, qPCR was performed to investigate the expression of the *sodA* gene in strain 654Δ*rnc* (**Figure 5b**). The level of mRNA transcript of *sodA* in <u>rnc</u> mutant strain was much higher than that in the parental strain *S.* Enteritidis 654. On the other hand, the level of *sodA* mRNA transcript in 654Δ*rnc*/p-*rnc* was similar to that of WT strain *S*. Enteritidis 654. These findings therefore showed that the absence of *rnc* gene actually resulted in an increase of *sodA* mRNA level in *S*. Enteritidis. To further test the level of production of SodA protein in strains carrying various constructs, Western blot was performed on the total cell lysates of these strains (**Figure 5c**). The result showed that production of SodA protein was actually reduced in the 654Δ*rnc* strain when compared to the parental wildtype strain 654; hence this finding was inconsistent with the observation that this strain produced a large amount of the *sodA* mRNA transcript. On the other hand, the quantity of SodA detectable in strain 654Δ*rnc*/p-*rnc* was similar to that of 654. Taken together, we found that, although the deletion of *rnc* gene caused an increase in the amount of mRNA transcript of the *sodA* gene in *Salmonella*, the production of SodA protein was reduced. This apparent paradox could be due to the stabilization of paired antisense and sense RNAs from the *sodA* gene resulting in the inhibition of *sodA* mRNA translation in 654Δ*rnc* mutant. *sodA* gene knockout mutant 654Δ*sodA* produced an ROS level similarly to that produced in 654Δ*rnc*, which was significantly higher than that in 654 and 654Δ*rnc*/p-*rnc* (**Figure 5d**). Based on these findings, we conclude that the *rnc* gene played an important role in mediating the translation of SodA protein, resulting in the regulation of ROS metabolism, in *S*. Enteritidis. ROS is a major weapon utilized by phagocytes to destroy internalized pathogens. A reduction in the ability to neutralize ROS in *Salmonella* could result in severe impairment in bacterial virulence (11).

**Figure 5.**
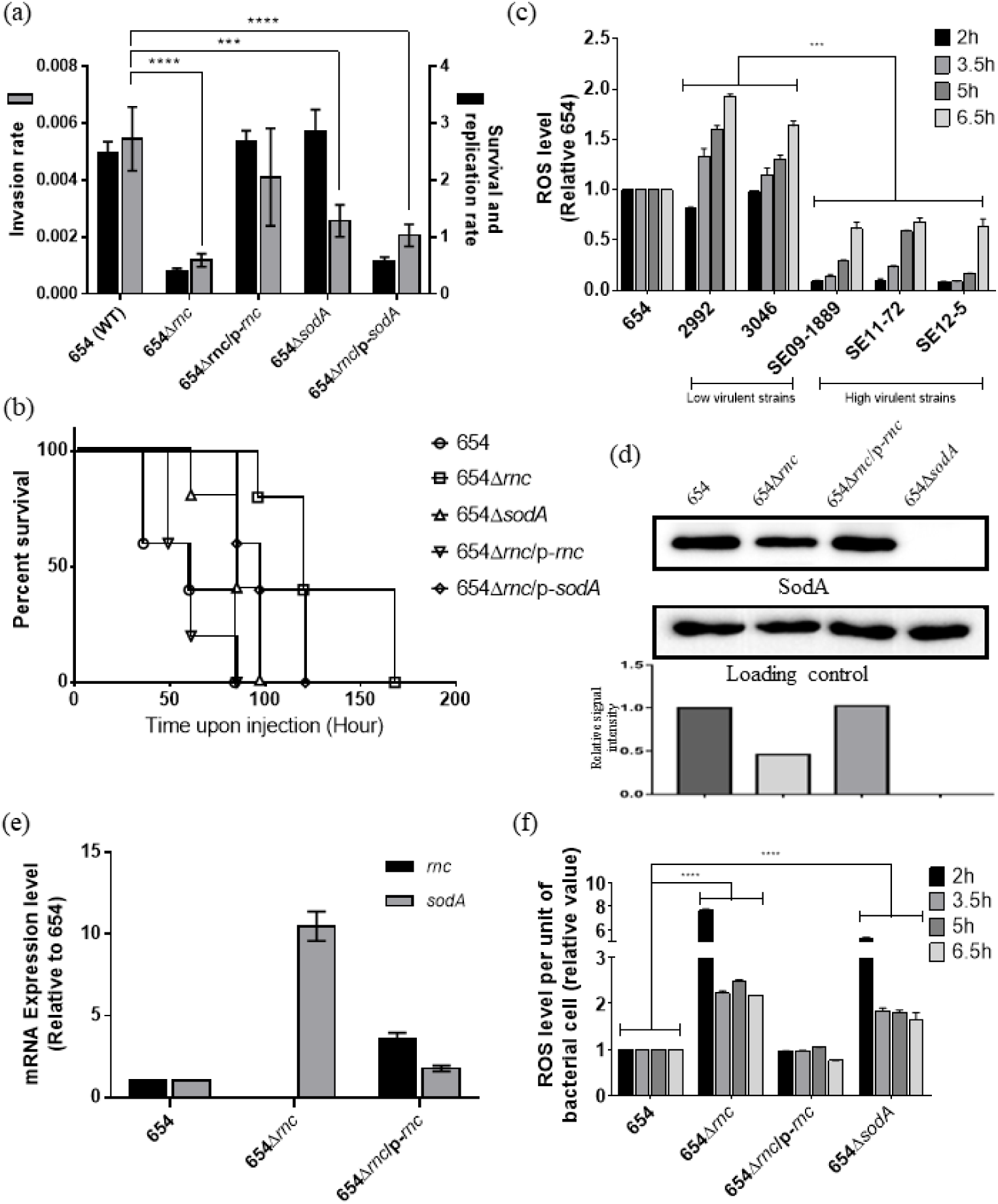
Virulence assays of the high virulence food isolate 654 and the corresponding *rnc* gene knockout mutant 654Δ*rnc*. (a)Macrophage internalization and replication assay. (b)Murine sepsis infection assay. The vector pET-*rnc* which carried the *rnc* gene was transformed into strain 654Δ*rnc* to produce 654Δ*rnc*::pET-*rnc,* which was then used in a complementation test. (c)ROS level in *Salmonella* isolates at different time intervals upon treatment with H_2_O_2_. (d)Western blot of the SodA protein in strains carrying different constructs. GAPDH was used as endogenous control. (e)The mRNA transcript level of the *sodA* gene in strains carrying different constructs. (f) The level of ROS in strains carrying different constructs.

To further confirm that the reduced virulence of strain 654Δ*rnc* was due to the reduced production of SodA, the p-*sodA* vector was transformed into strain 654Δ*rnc* to produce 654Δ*rnc*/p-*sodA*. The *sodA* gene in the vector was over-expressed to produce SodA protein. The RAW264.7 cell invasion and survival and replication assay were carried out to determine the virulence level of strains carrying various constructs (**Figure 5e**). Deletion of the *rnc* gene from *S*. *Enteritidis* 654 was found to dramatically lower the invasiveness, as well as the survival fitness and replication rate to level about 1/5 of that of the wild-type strain. The deletion of the *sodA* gene did not affect invasiveness of *Sallmonella* into RAW264.7 cels, while causing a 50% reduction of survival and replication rate. These results confirm that SodA plays an important role in the survival of *Salmonella* in the macrophage. The transformation of a *rnc*-bearing plasmid to the knockout mutant 654Δ*rnc* restored its intracellular survival and replication rate similarly to wild-type strain. The intracellular survival and replication rate of strain 654Δ*rnc/*p-*sodA* was about 3-fold that of 654Δ*rnc*, suggesting that introduction of the *sodA* gene can partially restore the function of *rnc*. Similar results were observed in mouse infection assay (**Figure 5f**), in which the virulence level of strain 654Δ*rnc*/p-*sodA* was comparable to the parental strain 654. These findings further confirmed that reduction in the virulence level of strain 654Δ*rnc* could be at least partly due to the lowered level of SodA protein.

## DISCUSSION

Antisense RNA (asRNA) is known to ubiquitously exist in bacterial genome and can be transcribed from the negative strands of protein-encoding genes. This type of RNA exhibits perfect complementation to RNA transcripts of the sense strand of the gene and form double-stranded RNAs (dsRNAs) by base-pairing with the mRNAs to regulate expression of specific genes (12). RNase III, a double-stranded RNA (dsRNA)-specific riboendonuclease encoded by the *rnc* gene, is implicated in cleavage of dsRNAs (13). The molecular mechanisms and biological significance of asRNAs in bacteria are still poorly defined. The widespread presence of asRNAs in bacteria could be transcription noises which are rapidly degraded by RNases; alternatively, these molecules may play a role as gene regulators that fine-tune the expression level of specific genes. Recently, cellular activities that determine the degree of accumulation of dsRNA have been reported to be involved in regulation of both bacterial infectivity and host immune responses (14, 15). Currently available data also suggest that the functional roles of RNase III in *Salmonella* involve regulation of the level of specific asRNA to control bacterial gene expression; such function, which may be mediated through non-specific regulation of the dsRNA level, may indirectly cause changes in the virulence level of *Salmonella*. We therefore investigated the mechanisms of RNase III in regulating *Salmonella* virulence.

In this study, we first observed that the *rnc* gene which encodes RNase III was over-expressed in the high virulence strains of *Salmonella*. We generated the *rnc* knockout mutants to investigate its role in expression of the virulence phenotype and found that the Δ*rnc* mutants indeed exhibited decreased intracellular survival and replication rate in macrophage cells (**Figure 2b**). Reduction in virulence level in *Salmonella,* when *rnc* was deleted, was confirmed by testing in murine sepsis model (**Figure 5b**). It is known that *Salmonella* is exposed to an abundance of ROSs produced by the host during infection, including superoxide (O^2−^), hydrogen peroxide (H_2_O_2_) and hydroxyl radical (OH^3−^) (16). These ROSs were important tools by which the immune system of the host utilizes to eradicate pathogens via exerting strong oxidative effects. It was reported in previous studies that the ability of *Salmonella* to neutralize ROS was important for its survival and propagation in the host, and that such function was mediated through production of the enzyme manganese superoxide dismutase (Mn-SOD) (17–19). Our results confirmed that the production of Mn-SOD in the Δ*rnc* mutant was indeed altered (**Figure 5d**). The transcription of *sodA* gene, encoding a kind of Mn-SOD, in the 654Δ*rnc* strain was significantly higher than its parental strain 654 but western blot showed that the production of SodA protein was significantly reduced in 654Δ*rnc*. These contradictory findings appear to suggest that deletion of *rnc* in *Salmonella* results in reduced RNase III production, which may cause accumulation of dsRNA including that of the transcription product of *sodA*, hence high transcript level of *sodA* was detectable. On the other hand, the asRNA of *sodA* could not be removed due to the lack of RNase III. Hence, the translation of *sodA* transcript would be inhibited, resulting in decreased production of SodA protein (**Figure 6a**). In this study, we performed an ROS assay and confirmed that the ROS content in *Salmonella* significantly increased when the *rnc* gene was deleted. This finding supports the idea that deletion of the *rnc* gene exhibits the effect of suppressing production of SodA protein production by inhibiting translation of the *sodA* transcript, hence, indirectly affect survival of *Salmonella* in the host cells.

**Figure 6.**
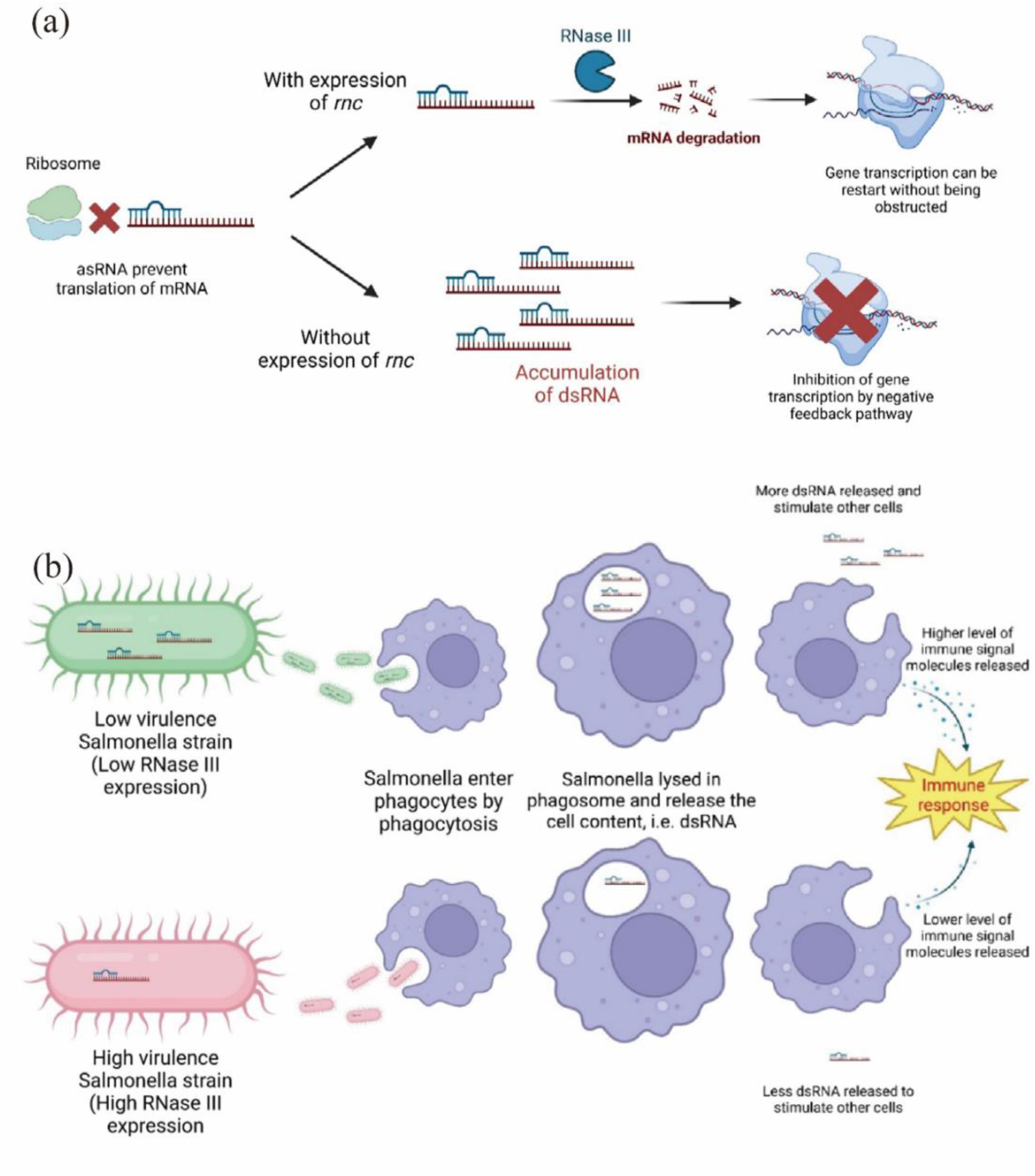
Mechanism underlying the contributions of RNase III to *Salmonella* virulence. (a) Demonstration of gene expression inhibition due to the accumulation of dsRNA. (b) Illustration of how host immune response triggered by dsRNA from low virulent and high virulent *Salmonella* strains.

Apart from demonstrating the role of RNase III in regulating the expression of the SodA protein, we also tested the role of RNAse III in regulating the host immune response to RNAs of bacterial origin. Firstly, we found that the RNAs extracted from the *rnc* knockout mutant indeed induced significantly higher levels of immune responses in the host, including the innate immune factors IFN-β, RIG-I and MDA-5, when compared to RNAs from the wild-type strain. Secondly, we showed that dsRNAs were only detectable in *rnc* knockout mutant (**Figure 1b**), suggesting that bacterial dsRNAs can be recognized by intracellular innate immune sensors of the cells and trigger immune responses in the host. The retinoic acid-inducible gene I (RIG-I) and melanoma differentiation-associated gene 5 (MDA-5) were known to be responsible for combating viral infection (20). Double stranded RNA is commonly found in RNA viruses as their genetic materials. The innate immunity elicited against dsRNAs plays an essential role in protecting human against various viral infections. Our experimental results showed that the dsRNAs in *Salmonella* could also trigger the activation of both RIG-I and MDA-5 pathways. A previous study showed that the alpha/beta interferon-based immune defenses could be triggered by RIG-I and MDA-5 (21). Our qPCR data showed that expression of IFN-β in RAW264.7 and Caco-2 cells significantly increased when the cells were transfected with RNA extracted from the *rnc* knockout mutant (**Figure 3b-e)**. Importantly, testing the effect of treatment by RNase III and Exonuclease T enzymes confirmed that such response was only triggered by bacterial dsRNAs but not ssRNAs (**Figure 4e)**. We therefore conclude that the dsRNA from *Salmonella* may act in a manner similar to that of viral dsRNAs by triggering the RIG-I and MDA-5 pathways in the host. Particularly, the generation of interferons in non-phagocytes may serve as the defense signal for the host to restrict the spreading of intracellular bacteria, like *Salmonella*. Over-expression of RNase III in the high virulence *Salmonella* strains could therefore remove excess dsRNAs and help the bacteria to evade the host immunity (**Fig 6b**).

To conclude, findings in this work suggest that RNase III played an important role in regulating the expression of Mn-SOD and bacterial resistance to oxidative stress which is important for survival of *Salmonella* in the host during the infection process. On the other hand, we also found that dsRNA was responsible for induction of the host immune response which is suppressed by RNase III. Over-expression of RNase III in *Salmonella* may result in a lower dsRNA level, leading to a milder innate immune response inducible by *Salmonella*. These novel findings pave the way to devise effective approaches to attenuate bacterial virulence by suppressing specific housekeeping and stress response genes including RNase III, which degrades dsRNA, a key factor that triggers the host immune response.

## MATERIALS AND METHODS

### RNA extraction, qRT-PCR and Immunoblotting

The overnight culture of the test strains was first re-inoculated into fresh LB broth and allowed to grow at 37℃ with shaking, until optical density reached 0.5; 1 mL of log-phase culture was then treated with the QIAGEN RNAprotect Bacterial Reagent. Total RNA was extracted by the Qiagen RNeasy Bacteria Minikit, followed by DNase treatment. For RNA extraction of RAW264.7 and Caco-2 cells, the cultured cells were treated with the QIAGEN RNAprotect Cell Reagent, followed by extraction of RNA by the Qiagen RNeasy Cell Minikit. The quality and quantity of RNA was determined by using the Nanodrop spectrophotometer. 1 μg of total RNA was subjected to reverse transcription using Life technologies Superscript III reverse-transcriptase. Real-time RT-PCR was performed by using the Applied Biosystem Quant Studio 3 and the Life Technologies SYBR Select Master mix. Melt curve analysis of PCR product was performed to ensure specificity of the selected primers. Expression levels of the test genes were normalized with a housekeeping gene that encodes the DNA gyrase subunit B for bacterial samples or GAPDH protein for cell samples. Primers used in qPCR are listed in **Table S1**. For the RNA immunoblotting assay, bacterial total RNA was quantified and an equal amount of RNA was separated on agarose gel, followed by transfer to PVDF membrane and detection using J2 antibody (Abcam #ab288755).

### Macrophage invasion and survival assay

The virulence level of the tested *Salmonella* strains was determined by infecting RAW 264.7 cells and measuring the internalization and replication rate. The bacterial strains were inoculated into fresh LB broth and incubated at 37℃ with shaking, until the optical density at 600 nm (OD600) of the bacterial cultures reached 0.5. The bacterial cells were harvested by centrifugation and washed once with phosphate buffered saline (PBS). The washed bacterial cells were then resuspended in DMEM cell culture medium, followed by addition to RAW 264.7 (ATCC® TIB­71™) cells pre-coated in a 24-well cell culture plate with a multiplicity of infection (MOI) ratio of 10:1. The plates were then centrifuged at 500 rpm for 5 min to synchronize the infection, followed by incubation at 37C, 5% CO_2_ for 25 mins. The medium was then removed and replaced by DMEM supplemented with 200 μg/ml gentamicin, and subjected to incubation for 1.5 h; DMEM containing 10 μg/ml of gentamicin was then used for the rest of the experiment. The supernatant was removed at 2 and 16 h after infection, the cells were then washed twice with pre-warmed PBS and lysed with 0.2% Triton X-100. Serial dilutions of the lysates (10^0^, 10^-1^,10^-2^,10^-3^,10^-4^) were plated onto LB agar to enumerate the intracellular bacteria.

### ROS assay

Overnight cultures of the test *Salmonella* strains were diluted in LB broth at OD600 of 0.08 and incubated at 37 °C with aeration until OD600 reached 0.3, followed by addition of 10 mM H_2_O_2_. 1 ml culture aliquots were collected at 2, 3.5, 5 and 6.5 h upon the addition of H_2_O_2_. The bacterial cells were pelleted and washed three times with phosphate-buffered saline (PBS), and resuspended in 500 µl of PBS containing 5 µM CM-H2DCFDA after the final wash, followed by incubation at 37°C in dark for 30 min. The labeled cells were washed once with PBS and resuspended in 500 µl of PBS. The fluorescence signal of a 200 µl aliquot was measured by Molecular Devices SpectraMax iD3 at 485±5nm excitation and 535±5nm emission wavelengths. The results were normalized according to the viable cell counts. Briefly, 100 µl were removed and subjected to 10-fold dilution. 100 µl from various dilutions were spread onto LB agar and incubated at 37 °C until colony forming units (CFUs) formed. CFU was counted in the various dilutions which produced distinct and countable colonies (25-250 CFUs).

### Gene knockout and cloning experiments

Gene knockout mutants of *Salmonella Enteritidis* strain 654 were generated by the λ-red system, following the protocol described by Ruth, et al [11]. The pKD46 plasmid was used as the helper plasmid and pKD4 plasmid was used as the template of kanamycin resistance gene. The mutants produced and the sequence of primers used are listed in **Table S1**. PCR was performed by using the high-fidelity polymerase to ensure the integrity of gene sequences. The voltage of electroporation was 18kV/cm. The mutants were recovered by addition of pre-warmed SOC medium and incubated at 37℃ with shaking for 2 hours. The recovered cells were selected on agar plates supplemented with kanamycin (50 μg/mL). Deletion of the target gene was confirmed by Sanger sequencing.

Gene cloning was performed by using the double restriction digestion method. Briefly, the pET-28b plasmid was chosen as the vector of desired gene sequences. The desired genes were amplified by high-fidelity PCR. Both the vector and the PCR product were digested by restriction enzymes at 37℃ for 4 hours, followed by ligation at 16℃ overnight, using T4 ligase. The ligation products were transformed into *E*. *coli* strain DH5α, followed by selection of transformants which had acquired the cloned gene. The genetic sequence of the selected clones was confirmed by Sanger sequencing. The recombinant plasmid harbored by the selected clones was then extracted and transformed into the target strains. The sequence of primers used are listed in **Table S1**.

### Murine sepsis infection model

ICR mice aged 5 weeks were used as the host for *S*. Enteritidis infection. Each experimental group consisting of 5 mice was infected by different mutants of *S*. Enteritidis. Briefly, *S*. Enteritidis mutants were first grown in LB broth until the optical density at 600nm reached 0.5. The bacterial cells were harvested and washed once with sterile 0.9% sodium chloride solution. The washed bacterial cells were then inoculated into 0.9% sodium chloride solution. The bacterial suspensions were injected into the mice through tail vein at the final dosage of 10^5^ bacterial cells. Water and food were given to each mouse during the experiment. The death rate of the mice in each experimental group was recorded at 12-hour intervals.

### Western blotting of SodA

*Salmonella* strains which harbored the desire constructs were first streaked on LB agar plate to ensure no contamination of the stock. The single colonies of each test strain were inoculated into LB broth and incubated overnight at 37℃ with shaking. The culture was re-inoculated to fresh LB broth and incubated at 37℃ with shaking until OD600 reached 0.5. The bacterial cells in 1 ml culture were pelleted by centrifugation and resuspended in 400 μL SDS loading buffer, then boiled for 10 minutes. Solubilized proteins were separated by SDS-PAGE and subsequently transferred to the PVDF membrane through Bio-Rad Trans-Blot Turbo Transfer System. Western blotting was carried out by probing the membrane with rabbit anti-SodA polyclonal antibody, followed by goat anti-rabbit IgG. The signal was developed by the addition of HRP-substrate and visualized by Bio-Rad ChemiDoc Imaging System. *Salmonella* GAPDH-specific antibody was used as endogenous loading control.

### Transfection of RNA

The corresponding cell line was cultured in a six-well plate, with cell density of 10^6^ cells per well. A total of 2.5 μg RNA was used for the transfection process of each well. Briefly, total RNA extracted from *Salmonella* strains carrying the constructs was first diluted with the Opti-MEM reagent (Gibco^TM^ #A4124801) and mixed with the Lipofectamine 2000 transfection reagent (Invitrogen^TM^ # 11668019) to form RNA-lipid complexes. The complexes were then added to active cell cultured in DMEM. The transfection process was allowed to proceed overnight at 37℃ under a supply of 5% CO_2_.

### Statistical analysis

All data were presented as the mean ± SD. One-way ANOVA analysis of variance was used to calculate the differences between various experimental groups. A two-tailed value of P < 0.05 was regarded as statistically significant. *P ≤ 0.05, **P ≤ 0.01, ***P ≤ 0.001, ****P≤0.0001.

## DECLARATIONS

### Availability of data and material

All datasets analyzed in the present study are available from the corresponding author on reasonable request.

## ACKNOWLEDGEMENTS

This study was funded by the Theme Based Research Scheme (T11-104/22-R), the Research Impact Fund (R1011-23) and Postdoctoral Fellowship (PDFS2223-1S09) from the Research Grant Council of the Government of Hong Kong SAR. L.H. was supported by Wang-Cai Biochemistry Lab donation grant (21KCNGO016) and a Kunshan Double Innovation Talents Award (KSSC202102069).

